# NSeqVerify: An Easy-to-Use Desktop Suite for Integrated NGS Data Analysis, from Raw Reads to Taxonomic Assignment

**DOI:** 10.1101/2025.10.31.685854

**Authors:** Roberto Reinosa Fernández

**Affiliations:** HIV-1 Molecular Epidemiology Laboratory, Microbiology Department, Ramon y Cajal University Hospital-Ramon y Cajal Institute for Health Research (IRYCIS), Madrid, Spain

## Abstract

**Motivation:** The proliferation of next-generation sequencing (NGS) data has created a computational bottleneck, especially for researchers lacking specialized bioinformatics training. Standard analysis workflows require mastering multiple command-line tools, hindering exploratory data analysis and delaying scientific discovery.

**Results:** This work presents **NSeqVerify**, a new cross-platform, open-source desktop software developed in Java, designed to overcome these barriers. NSeqVerify implements a fully integrated genomic workflow within a single, intuitive graphical user interface (GUI). The suite includes: (1) a **preprocessing module** for quality control and filtering of FASTQ files; (2) a **de novo assembler** employing a sophisticated De Bruijn graph algorithm with an iterative multi-k-mer strategy to maximize contiguity; and (3) a **taxonomic assignment module** that automates BLAST searches against NCBI databases and displays the results in an easily interpretable tabular format. The tool was validated through controlled use cases, demonstrating its ability to accurately reconstruct reference viral genomes (HIV-1) and to deconvolute metagenomic mixtures (HIV-1 and SARS-CoV-2). The final test consisted of analyzing a real elephant fecal virome (SRA: SRR35776009), where NSeqVerify successfully assembled contigs — two of which overlapped and appeared to form a partial 1555 bp genome of a putative *Smacovirus*, enabling the identification of its capsid protein and the prediction of its 3D structure using AlphaFold.

**Conclusion and Availability:** NSeqVerify democratizes NGS data analysis, providing a robust “all-in-one” solution that empowers molecular biologists, students, and clinicians to perform end-to-end genomic analyses. The software is freely available under the GNU GPLv3 license at (https://github.com/roberto117343/NSeqVerify).

## 1. Introduction

Next-generation sequencing (NGS) has revolutionized the life sciences, enabling unprecedented-scale research in fields such as metagenomics, virology, clinical genomics, and evolutionary biology [1]. However, the resulting “data deluge” has shifted the main challenge from data generation to analysis and interpretation. A typical short-read genome analysis workflow is a multi-step process requiring substantial computational expertise.

This canonical process begins with rigorous quality control (QC) and preprocessing of raw data (FASTQ format), using tools such as **FastQC** [2] and **Trimmomatic** [3]. Next, the filtered reads are **assembled de novo** to reconstruct the original genome — a computationally intensive task for which sophisticated algorithms have been developed, such as those used by **SPAdes** [4], **MEGAHIT** [5], or **Velvet** [6]. Finally, the resulting contigs must be **annotated** to determine their taxonomic origin and potential function, often involving large-scale homology searches using **BLAST** [7] against public databases. Each of these steps not only requires a different command-line tool but also dependency management, format conversion, and parameter optimization.

This “command-line barrier” represents a significant obstacle for the research community. To bridge this gap, **NSeqVerify** was developed based on three guiding principles: (1) **Integration**, combining critical stages into a coherent workflow; (2) **Accessibility**, providing an intuitive GUI; and (3) **Reproducibility**, allowing users to easily document the parameters used.

## 2. System and Methods

**NSeqVerify** is a standalone desktop software built in Java, using the **Swing** library for its interface. This design ensures portability across **Windows, macOS**, and **Linux**.

### 2.2.1. Module 1: FASTQ Preprocessing (Preprocess FASTQ)

This module is the first step toward ensuring a high-quality assembly. It accepts a FASTQ file and performs user-configurable operations such as **Phred quality filtering, end trimming, reverse complement generation**, and **read subsampling**.

### 2.2.2. Module 2: De Novo Assembly (Assemble)

The core of NSeqVerify is its **assembler**, which implements a **De Bruijn graph (DBG)** algorithm [8]. Its key features include:

- **Multi-K-mer Strategy::** Assembly is performed iteratively using a series of user-defined k-mer sizes. The use of multiple k-mers has been shown to significantly improve assembly contiguity, especially in metagenomic data [9]. The contigs from one round are used as “super-reads” in the next.
- **Advanced Graph Simplification:** Heuristics are applied to prune “tips” and resolve “bubbles” — graph topologies commonly arising from sequencing errors and genomic variants [10].

### 2.2.3 Module 3: Taxonomic Assignment (Classify nt /Classify aa)

This final module provides an initial biological characterization of the contigs. It automates the submission of each contig to the NCBI web API for **BLASTn** or **BLASTp** searches [7]. The results are automatically parsed to extract key metrics from the best hit and are presented in a **tab-delimited text file (TSV)**. Contigs with no significant matches are isolated, highlighting them as candidates for further investigation.

## 3. Results and Validation

The performance and accuracy of NSeqVerify were evaluated through a series of rigorous case studies using both simulated and real metagenomic data.

### 3.1 Case Study 1: Accuracy in Reconstructing a Reference Genome (HIV-1)

To assess the assembler’s accuracy, 100,000 fragments of 150 bp were generated *in silico*from the HIV-1 reference genome (HXB2). Using a multi-k-mer strategy with odd k-mer values from 21 to 91, a minimum k-mer frequency of 5, and a minimum contig length of 200 bp, NSeqVerify reconstructed a pair of contigs, with the main one covering most of the reference genome. The assembly was nearly perfect, except for the terminal regions.

### 3.2 Case Study 2: Resolution of a Mixed Viral Metagenome

To test the tool’s ability to deconvolute complex data, a synthetic metagenome was created by mixing 75,000 150 bp reads from the **SARS-CoV-2** genome with 75,000 reads from **HIV-1**. The same parameters were used as in the previous case, except for the k-mers (odd lengths 21–81). NSeqVerify successfully separated the mixture, producing two distinct long contigs. Each contig corresponded to one of the original viral genomes, demonstrating the algorithm’s effectiveness in resolving and assembling individual genomes from a mixed sample. As in the previous case, the genomes were complete except for the terminal ends.

### 3.3 Case Study 3: Detection of a Viral Genome in a Real Virome

To validate NSeqVerify in a realistic discovery scenario, a public dataset from an elephant fecal virome (SRA: **SRR35776009**) was analyzed. The data (Illumina NovaSeq 6000, 250 bp paired-end) were preprocessed using NSeqVerify (*trim ends:*30, *max reads:*50,000, *reverse complement:*enabled). De novo assembly was performed using a **multi-k-mer strategy** (odd k-mers: 21–91), a minimum k-mer frequency of 5, and a minimum contig length of 200 bp. For filtering, the minimum quality was set to 0 since this value was not available in the FASTQ file.

This analysis produced numerous contigs. Among them, two overlapping contigs — **Contig_4 (1**,**471 bp)** and **Contig_5 (1**,**421 bp)** — showed nucleotide-level homology to viruses of the **Smacoviridae** family. Given their overlap, a final **consensus sequence of 1**,**555 bp** was generated using the **EpiMolBio** tool [12].

The **open reading frames (ORFs)** of this consensus sequence were analyzed using **OrfViralScan** [13]. The resulting protein sequences were classified with the **BLASTp** module of NSeqVerify, revealing a strong homology for one of the ORFs with **Smacovirus capsid proteins** (*ORF_6, 334 aa*). The closest hit was the capsid protein of *Smacoviridae sp*. (Accession: WCR62194.1), showing **51.66% amino acid identity** and an **E-value of 1e-102** over 90% of the sequence. The low identity percentages with the nearest relatives (48–55%) strongly suggest that this contig may represent a member of a **new viral species** within the *Smacoviridae*family.

This workflow — from the initial assembly to manual merging and final characterization — demonstrates the potential of NSeqVerify to produce high-quality contigs that can serve as the foundation for potential genomic discoveries in complex metagenomic samples. The partial genome sequence and its ORF are presented in **Appendix A**.

## 4. Discussion

NSeqVerify stands as a robust and validated solution for NGS data analysis, focused on **accessibility**. The goal is not to replace high-performance command-line tools, which will remain the gold standard for large-scale analyses on computational clusters. Instead, NSeqVerify fills a crucial niche: **accessible desktop analysis**.

When compared to the existing bioinformatics tool ecosystem, its main competitive advantage lies in its **vertical integration within a GUI**. While other graphical suites offer a broader range of features, they are often more complex and not specifically focused on this workflow. NSeqVerify specializes in guiding the user **from raw FASTQ files to classified contigs** in the most straightforward way possible.

### 4.1. Limitations and Future Directions

To provide a transparent evaluation of the tool, it is essential to acknowledge its current limitations, which define the path for future development. It should be noted that NSeqVerify is currently in an **alpha version**, but given its present utility, the release of the tool is more than justified.

1. **Scalability and Performance:** De novo assembly is a memory-intensive process. NSeqVerify is currently limited to processing **tens of thousands of reads**.
2. **Dependence on the BLAST Web Service:** The classification module relies exclusively on the **NCBI web service**, making it slow for thousands of contigs and requiring an internet connection.
3. **Handling of Paired-End Read Data:** The current workflow does not explicitly use paired-end read information, thereby losing valuable spatial data for **scaffolding**.
4. **Lack of Command-Line Interface (CLI):** The GUI-centered design limits **automation and large-scale reproducibility**.
5. **Exclusive Support for Illumina Reads:** The assembler is specifically designed for **short reads (Illumina technology)**.

In conclusion, **NSeqVerify** is a tool that significantly reduces the entry barrier to genomic analysis. By packaging powerful algorithms into an easy-to-use interface, it serves as an excellent **“first-look” and exploratory analysis tool**, enabling a broader community of scientists to obtain meaningful insights from their own data.

## 5. Availability

**NSeqVerify** is open-source software distributed under the terms of the **GNU General Public License v3**. It is available on GitHub at the following address: https://github.com/roberto117343/NSeqVerify

## 7. Figures and Appendices

**Figure 1.**
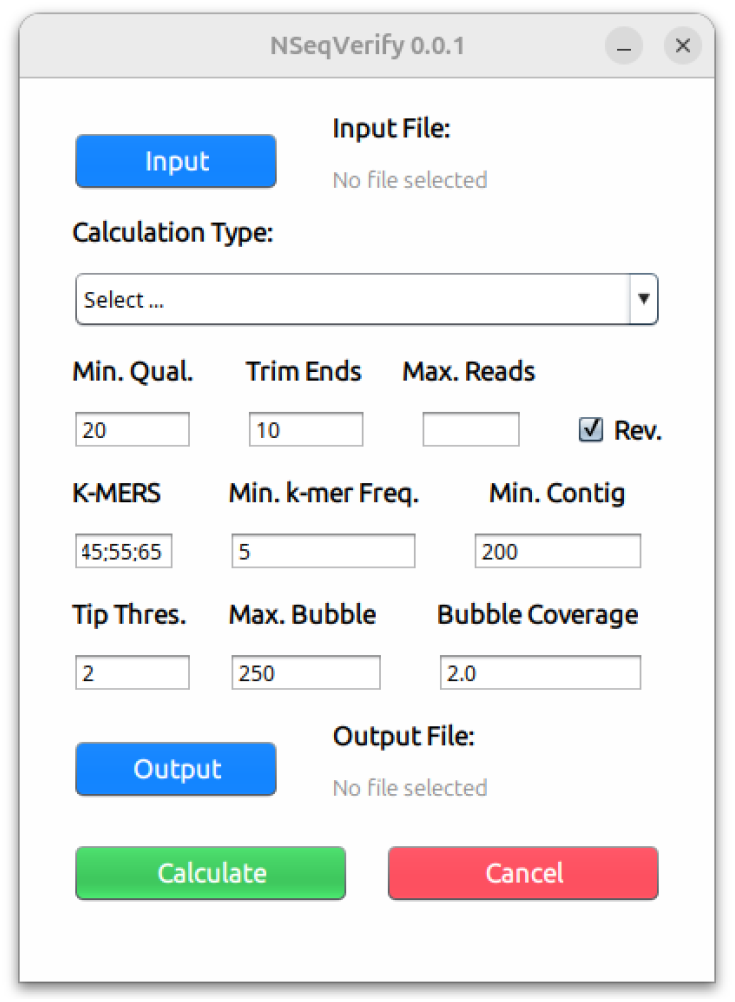
Graphical User Interface of NSeqVerify.

**Figure 2.**
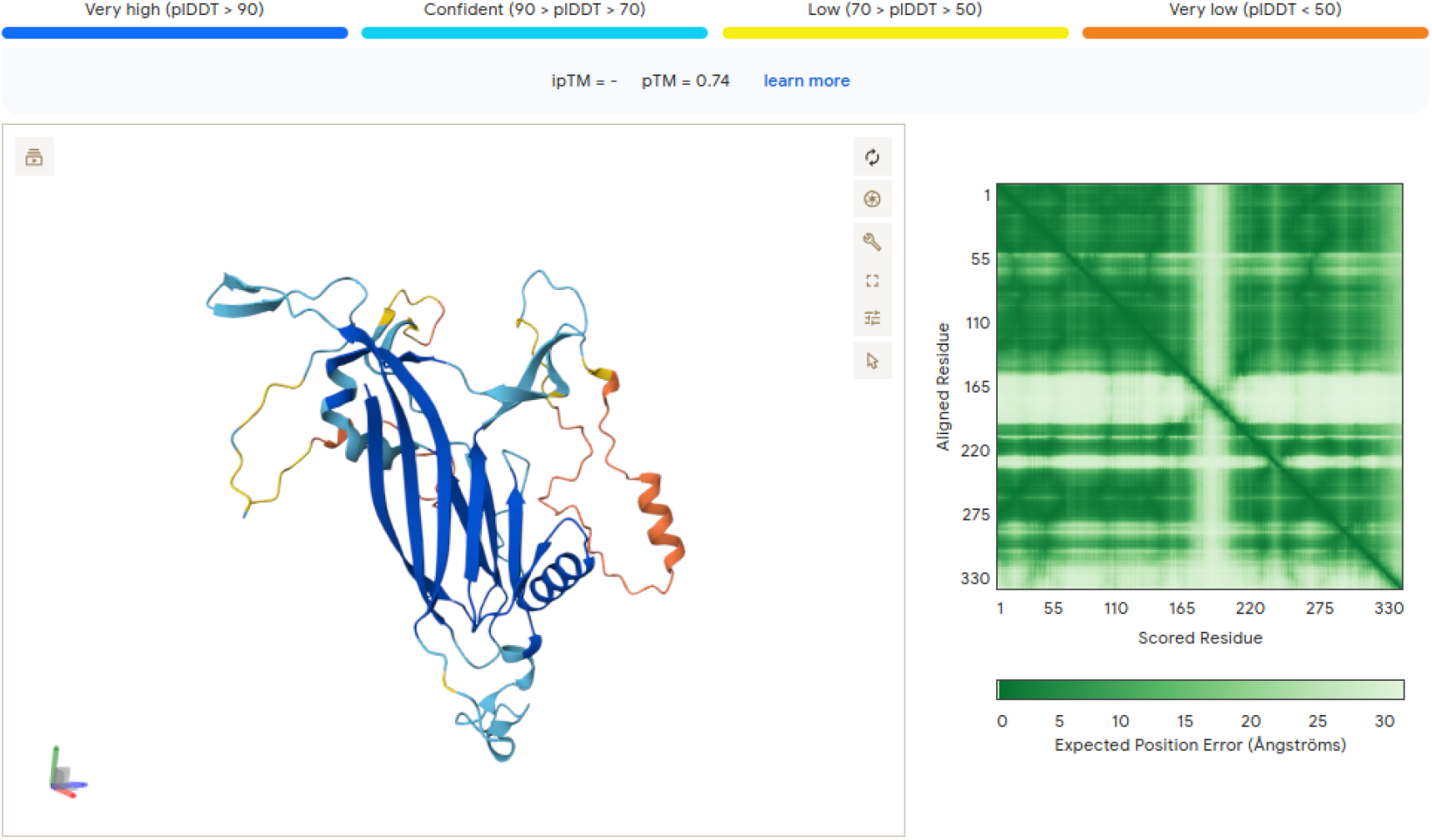
Predicted 3D Structure of the Putative *Smacovirus*Capsid Protein.

**Figure 3.**
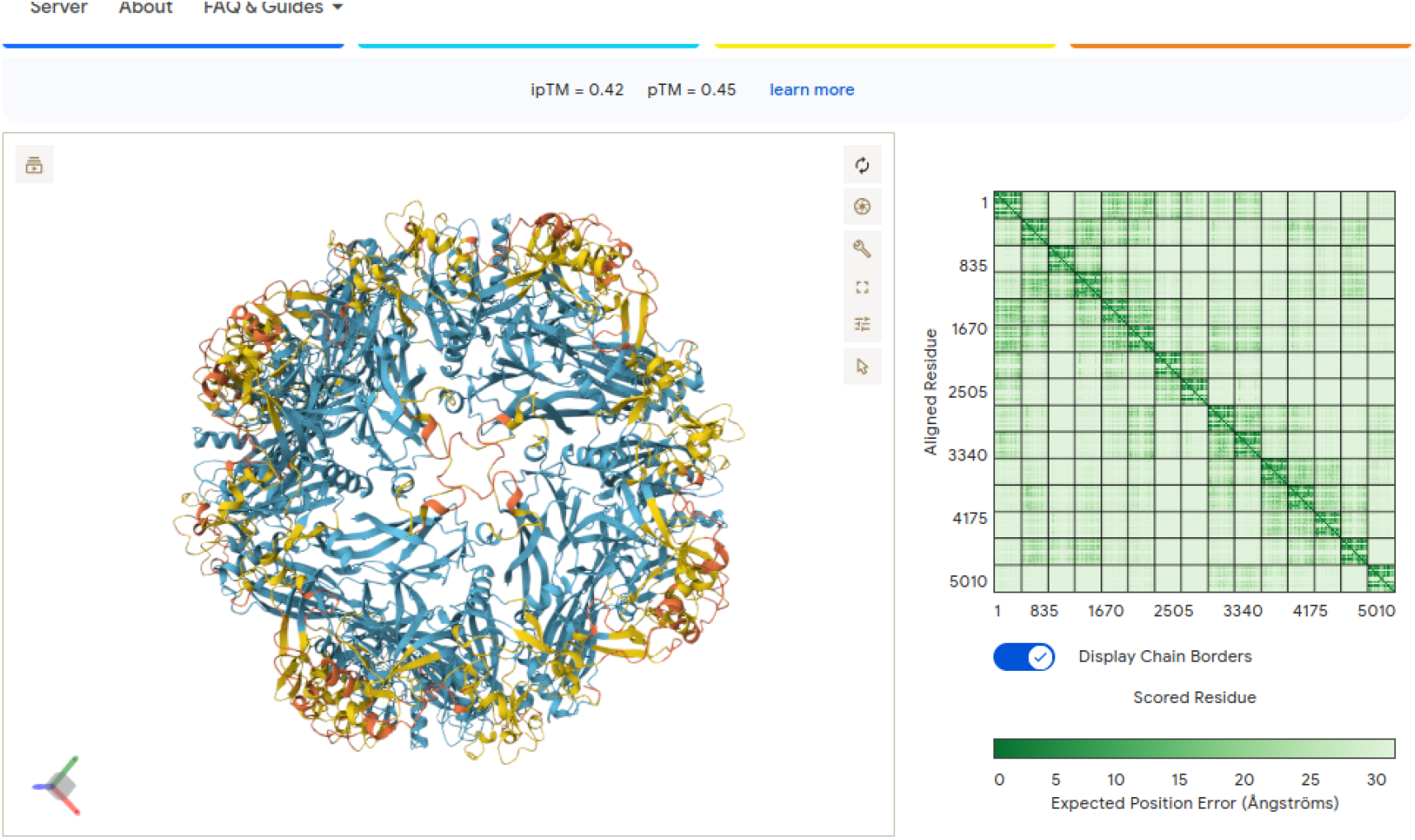
Predicted 3D Structure of the Putative *Smacovirus*Capsid Protein. Prediction of the assembly of 15 subunits (maximum allowed by AlphaFold), View 1.

**Figure 4.**
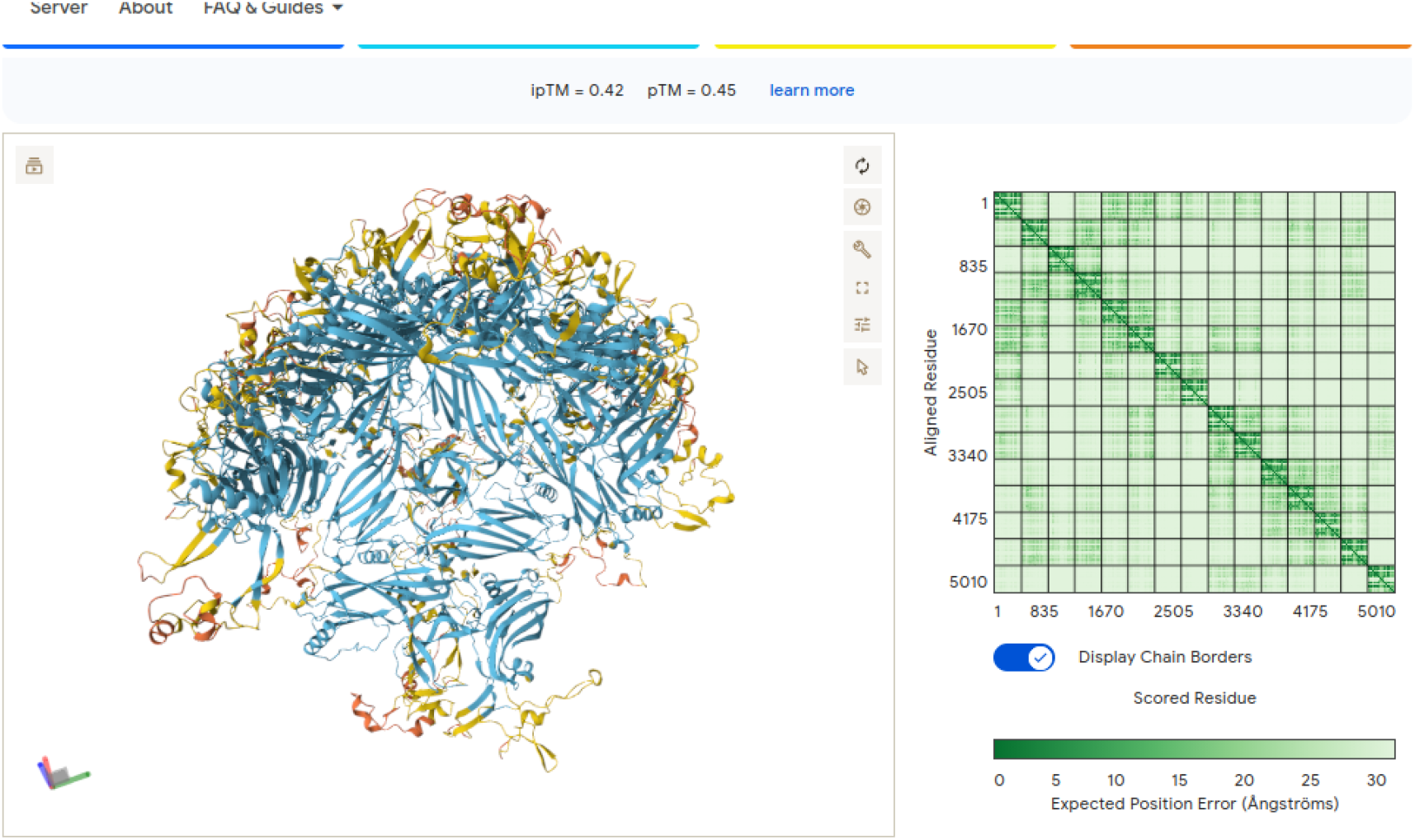
Predicted 3D Structure of the Putative *Smacovirus*Capsid Protein. Prediction of the assembly of 15 subunits (maximum allowed by AlphaFold), View 2.

## [APPENDIX A] Consensus Genome and ORF6 (Capsid Protein) Sequence of the Putative *Smacovirus*(1,555 bp)

### Putative Identification

*Partial genome of a presumably uncharacterized Smacovirus*.

### Contig Sequence (FASTA format)

>Smacovirus_consensus_length_1555

ACATATTCTTGCCTCTGTCAAGTTCTCGTCAAAGACGGTACAGCGCTTGACAATTATTAACGGAGGAACTA CTATGTATGGTTTTCGTCGTAGAAGGTATGGTTATCGTAGGAAGTCTAGGTATTCTAGGCGTAGGAGGTACTACTGATGGTTGTCCATCCTGTCCTGACGGCCGTAGGTCTTGCCGGTCTCGGAGTTTCCGCCGGTGCTAACGTCTACGCCCAGTATCGTCAGAGGCAGTTGTACCGCCAACAGGCTAATGCTTATTCTAACCTCCATCGTGGATACACAAAATATCTGAAATCCCATGGCAGACAAATCAACCCGGACCGCGCTTGGACGTCGTATTATGGCCAGTATCAGAGAGCATTGGCCAATTATGAGAGCAGTTATGCTGGTAGTTTTGGTACTGTCGGAGGTTCTGTCGGAGCTGGTTCAGCTATTGCGCAGCATTCTCTTAGATCCACGAATGGAACATTTAGGAGGTTACCCAGATGAAATACACATTTCAGCACTATATCGATATCAGTACCTCGGCTGAATCGATGCAGATCATTTCGGTTAATGCCGGTGGTCAGTATCTGATTAATCGTTGCAGACATCTTCTTGGAACTTACAAATACTACAAGCTCGGAAAGGTTTCCATTAGGCTCGTTCCGGCTTCTACTCTTCCTGTGGACCCTCTCGGCCTGTCCTATGCCGATACGGATCCGCAGACCGTCGACCCTCGCGACCAGCTTAATCCCGGTCTTGTCCGTATTACCAACGGTGAAGATTTCCAGTATTCGATTGATGGGGTCTCTAGTGCCTCACAGGACGAAATTTACAAAGCTATGATGCTCGACCCTCGTTGGTCTAAGTTCATGCTTCAGAGTGGCTTTAGGCGTTCAGCATCGCCCCTCTTCTGGTCCGTCGGTCAGCTCCACCAGGACGCATACCCCGGTTCAACTGTCAACGTCTTCACTAAAGGTACTGGACTACCCCAGACTAATTCTTGGATGTTTTCTTCTACTCGTTCCGGTACTACTGATGTTTCTGCTGCAATGAAGAATCTCGGTTCCGAGGGCATTAGGGTTAACTGTCAGGATTCGGATCCTCACGGATTTTTCCAGACTGGTCACCGTCAGCGTATGTCTTGGTTACCTACCGATATGCTGCAGCAGTTTGCCGGTGGTTCTTCTATTGCTTCCACGGATATGTTTACAATGGCCGGCCTTAATCCCATCTTGGCCCCCAATATTATCACGTGCATCCTGCCTAGAGCCTACAAGACACTGTACTACTACCGTCTTTTCATTACTGAAACTGTCTACTTCAGCGGTATCAAGAATGTTGGACTTGGCATTGAGGAGGCTGAAAGCTTGTACGAATACAACGGACTTGATAACTTCACTAACCCGATGTTCCCGACTGGGGTTAATCCGGTAAACGGAATTCAGATCCTGAAGACTTATTCGGAGCTTGTTACTCCGCCTAATGACGGTGATTCGAATGACTGAAATCAAGATCTATACATCTCTCGGTGTAGTCACCGCCAAGCCTATGAGCTTGGTAG

>Smacovirus_consensus_length_1555_rev_compl

CTACCAAGCTCATAGGCTTGGCGGTGACTACACCGAGAGATGTATAGATCTTGATTTCAGTCATTCGAATCACCGTCATTAGGCGGAGTAACAAGCTCCGAATAAGTCTTCAGGATCTGAATTCCGTTTACCGGATTAACCCCAGTCGGGAACATCGGGTTAGTGAAGTTATCAAGTCCGTTGTATTCGTACAAGCTTTCAGCCTCCTCAATGCCAAGTCCAACATTCTTGATACCGCTGAAGTAGACAGTTTCAGTAATGAAAAGACGGTAGTAGTACAGTGTCTTGTAGGCTCTAGGCAGGATGCACGTGATAATATTGGGGGCCAAGATGGGATTAAGGCCGGCCATTGTAAACATATCCGTGGAAGCAATAGAAGAACCACCGGCAAACTGCTGCAGCATATCGGTAGGTAACCAAGACATACGCTGACGGTGACCAGTCTGGAAAAATCCGTGAGGATCCGAATCCTGACAGTTAACCCTAATGCCCTCGGAACCGAGATTCTTCATTGCAGCAGAAACATCAGTAGTACCGGAACGAGTAGAAGAAAACATCCAAGAATTAGTCTGGGGTAGTCCAGTACCTTTAGTGAAGACGTTGACAGTTGAACCGGGGTATGCGTCCTGGTGGAGCTGACCGACGGACCAGAAGAGGGGCGATGCTGAACGCCTAAAGCCACTCTGAAGCATGAACTTAGACCAACGAGGGTCGAGCATCATAGCTTTGTAAATTTCGTCCTGTGAGGCACTAGAGACCCCATCAATCGAATACTGGAAATCTTCACCGTTGGTAATACGGACAAGACCGGGATTAAGCTGGTCGCGAGGGTCGACGGTCTGCGGATCCGTATCGGCATAGGACAGGCCGAGAGGGTCCACAGGAAGAGTAGAAGCCGGAACGAGCCTAATGGAAACCTTTCCGAGCTTGTAGTATTTGTAAGTTCCAAGAAGATGTCTGCAACGATTAATCAGATACTGACCACCGGCATTAACCGAAATGATCTGCATCGATTCAGCCGAGGTACTGATATCGATATAGTGCTGAAATGTGTATTTCATCTGGGTAACCTCCTAAATGTTCCATTCGTGGATCTAAGAGAATGCTGCGCAATAGCTGAACCAGCTCCGACAGAACCTCCGACAGTACCAAAACTACCAGCATAACTGCTCTCATAATTGGCCAATGCTCTCTGATACTGGCCATAATACGACGTCCAAGCGCGGTCCGGGTTGATTTGTCTGCCATGGGATTTCAGATATTTTGTGTATCCACGATGGAGGTTAGAATAAGCATTAGCCTGTTGGCGGTACAACTGCCTCTGACGATACTGGGCGTAGACGTTAGCACCGGCGGAAACTCCGAGACCGGCAAGACCTACGGCCGTCAGGACAGGATGGACAACCATCAGTAGTACCTCCTACGCCTAGAATACCTAGACTTCCTACGATAACCATACCTTCTACGACGAAAACCATACATAGTAGTTCCTCCGTTAATAATTGTCAAGCGCTGTACCGTCTTTGACGAGAACTTGACAGAGGCAAGAATATGT

### ORF6 Sequence (FASTA format)

> ORF6 Capsid Protein

MKYTFQHYIDISTSAESMQIISVNAGGQYLINRCRHLLGTYKYYKLGKVSIRLVPASTLPVDPLGLSYADTDPQTVDPRDQLNPGLVRITNGEDFQYSIDGVSSASQDEIYKAMMLDPRWSKFMLQSGFRRSASPLFWSVGQLHQDAYPGSTVNVFTKGTGLPQTNSWMFSSTRSGTTDVSAAMKNLGSEGIRVNCQDSDPHGFFQTGHRQRMSWLPTDMLQQFAGGSSIASTDMFTMAGLNPILAPNIITCILPRAYKTLYYYRLFITETVYFSGIKNVGLGIEEAESLYEYNGLDNFTNPMFPTGVNPVNGIQILKTYSELVTPPNDGDSND

